# Differential equation based minimal model describing metabolic oscillations in *Bacillus subtilis* biofilms

**DOI:** 10.1101/775593

**Authors:** Ravindra Garde, Bashar Ibrahim, Ákos T. Kovács, Stefan Schuster

## Abstract

Biofilms offer an excellent example of ecological interaction among bacteria. Temporal and spatial oscillations in biofilms are an emerging topic. In this paper we describe the metabolic oscillations in *Bacillus subtilis* biofilms by applying the smallest theoretical chemical reaction system showing Hopf bifurcation proposed by Wilhelm and Heinrich in 1995. The system involves three differential equations and a single bilinear term. We perform computer simulations and a detailed analysis of the system including bifurcation analysis and quasi-steady-state approximation. We also discuss the feedback structure of the system and the correspondence of the simulations to biological observations. We also specifically select parameters that are more suitable for the biological scenario of biofilm oscillations. Our theoretical work suggests potential scenarios about the oscillatory behaviour of biofilms and also serves as an application of a previously described chemical oscillator to a biological system.

## Introduction

Development of a complex biofilm provides several benefits to bacteria, including efficient nutrient distribution, defence from chemical attacks, or in the case of a floating pellicle on the surface of liquids, better gaseous exchange. Biofilms are thus complex communities of bacteria and as such, many types of social dynamics come into play. One of these is the division of labour ^1,2^. The core of the biofilm growing on a solid surface shows a different metabolic state than the periphery. The periphery can freely access the nutrients from the surrounding environment. The interior, however faces hindrance in obtaining a stable inflow of nutrients because the peripheral cells use up the nutrients that diffuse towards the interior. An experimental setup to simulate that situation is provided by a microfluidics chamber ^1^.

An example of such a nutrient gradient is the production and diffusion of ammonia in the biofilm. Every cell in the biofilm has the ability to produce ammonia ^1,3^. However, this small chemical compound is highly diffusive and therefore escapes into the environment as soon as it is produced by the cells in the periphery, thus leading to waste of nitrogen. In the interior, the ammonia produced by the cells diffuses out into the periphery. Thus the interior cells monopolize ammonia production for the entire biofilm. Ammonia being an essential component of glutamine metabolism could be used to control the growth rate of the periphery by limiting its supply. The interplay between the inner and outer cells is required for glutamine synthesis and therefore the growth of the biofilm.

To understand biofilms more closely and make predictions based on empirical data, several models have been developed^1,4–8^. Liu *et al.*^1^ observed oscillations in the biofilm which they explained by different metabolic roles performed by the different compartments in the biofilm. They also established a model based on six differential equations. In our model, we have focused mainly on metabolism of glutamate among the various amino acids since glutamate and ammonia are both involved in the production of various amino acids through transamination, which are then equated to growth.

Since many biological oscillators have been described by less than six variables,^9,10^ a simpler model could be established for biofilm oscillations as well. Our ultimate aim was to develop a minimal model to describe the metabolic oscillations happening in a biofilm. Minimal models are the simplest way to describe a certain phenomenon with the least number of parameters and this is in agreement with Occam’s razor. For example, minimal models were established for glycolytic oscillations by Higgins^11^ and Selkov^12^ and for calcium oscillations by Somogyi and Stucki^13^.

Here we employ the smallest chemical reaction system showing Hopf bifurcation^14^, which was further analysed15,16 and used to describe p53 oscillations^17^. In particular, Wilhelm and Heinrich ^14,15^ performed a thorough stability analysis of the model. We test to what extent the terms in this model match the processes in a biofilm system. In this analysis we focus on the Hopf bifurcation, discuss the feedback structure and point out the correspondence of the simulations to biological observations.

To study the effect and possible benefit of oscillations, it is of interest to compute the average values of variables, as was done for several oscillators^18–23^. For linear differential equation systems showing oscillations (such as the system describing the harmonic pendulum), the average values equal the values at the marginally stable steady state. For nonlinear differential equation systems, the average values of oscillations often differ from the values at the unstable steady state surrounded by the oscillations. However, there are some types of nonlinear systems for which equality holds, for example, Lotka-Volterra systems of any dimension^19^. The equality property has also been proved for some models of calcium oscillations^21,24^ and the Higgins-Selkov oscillator^21^. Here, we probe the model employed for describing biofilm oscillations for the above-mentioned property.

## Materials and methods

### The model

Based on the scenario described by Liu *et al.*, the biofilm was separated into two compartments - the interior and the periphery^1^. We tested several published models of oscillating systems^10–12,25–27^ and found that the smallest chemical reaction system with Hopf bifurcation^14^ matched the biological setup.. The term chemical system mathematically means that only up to bilinear terms are involved.

The model includes five reactions (see Figure 1) and three species with variable concentrations. The general variables *X*, *Y* and *Z* from the Wilhelm and Heinrich model can be assigned for the biofilm system to: peripheral glutamate (*G*_*p*_), ammonia (*A*) and internal glutamate (*G*_*i*_), respectively. Based on mass-action kinetics, the reactions have been translated as follows into a system of ordinary differential equations (ODEs):

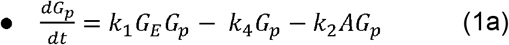

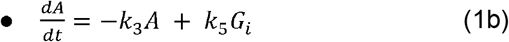

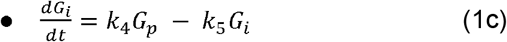

**Figure 1:**
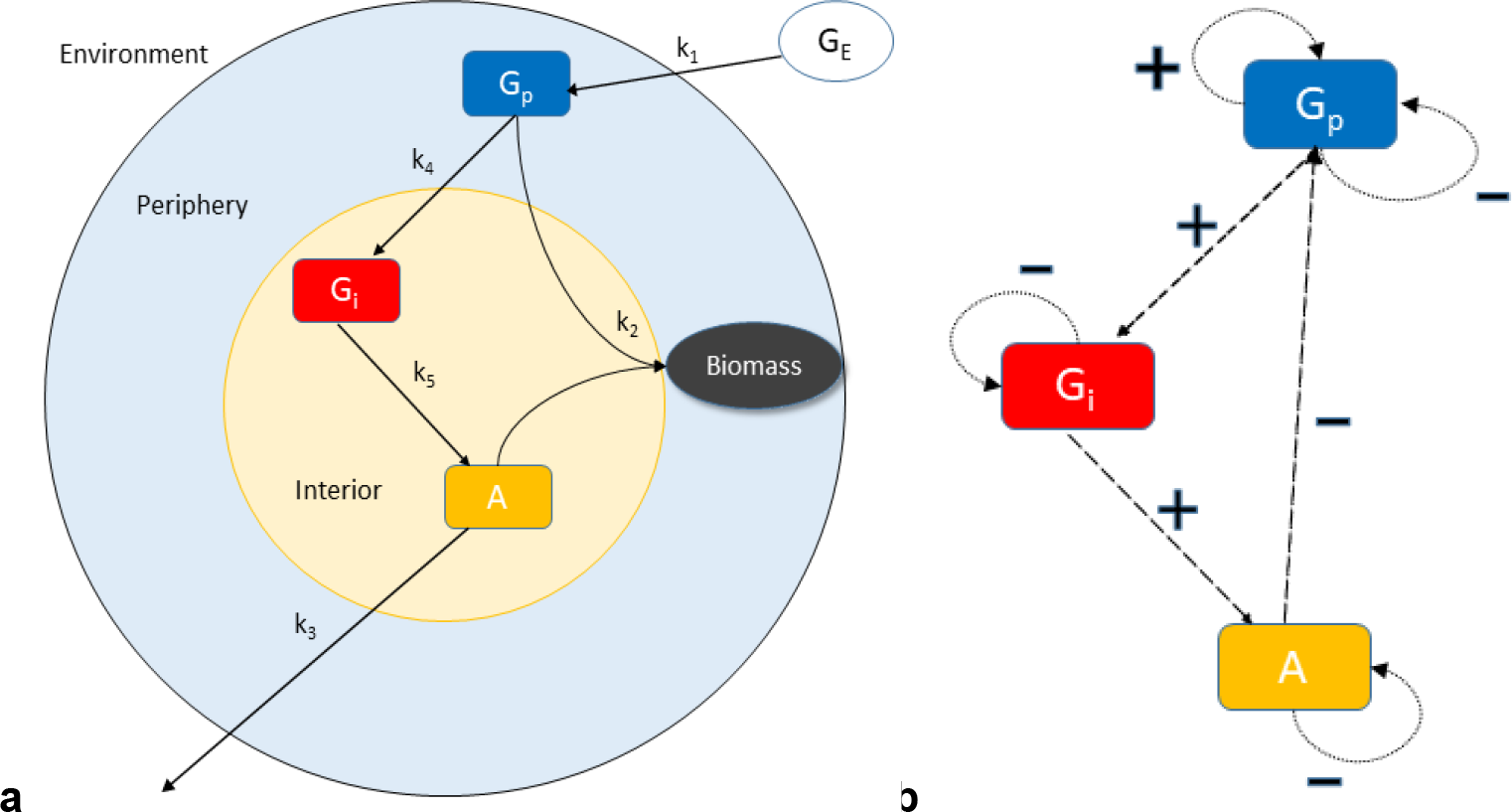
Schematic representation of the biofilm metabolic oscillation model. (**a**) The five reactions with rate constants *k*_*1*_ through *k*_*5*_ between the substances *G*_*i*_, *G*_*p*_, *A* having variable concentrations and *G*_*E*_ considered to be constant. The end result is the production of biomass. (**b**) Feedback structure of the model. *G*_*p*_ is self-amplifying while all three variables are self-degrading. *G*_*p*_ positively influences *G*_*i*_, which positively influences *A* which negatively influences *G*_*p*_, thus, the overall feedback is negative.

The terms in the equations are interpreted as follows:

- *k*_*1*_*G*_*E*_*G*_*p*_: Uptake of glutamate in the environment (*G*_*E*_) by the periphery of the biofilm
- *k*_*4*_*G*_*p*_: Diffusion of glutamate from the periphery of the biofilm into its interior
- *k*_*2*_*AG*_*p*_: Consumption of glutamate and ammonia to produce biomass.
- *k*_*5*_*G*_*i*_: Consumption of glutamate to produce ammonia
- *k*_*3*_*A*: Diffusion of ammonia into the surroundings.

### Simulation

For computer simulations, we used the software COPASI versions 4.16 and 4.24^28^ and its LSODA deterministic solver. The simulations were double-checked using the Matlab ode15s (MathWorks) function. The figures of the simulations were produced using COPASI, and the 3D phase plot was generated using Matlab plot3 function.

All the rate constants were adopted from the publication of Wilhelm and Heinrich^14^ and were rescaled such that the period of oscillations matched the one in the experimental work by Liu *et al.*^1^. The glutamate concentration in the environment was also adopted from the latter paper.

The predicted doubling time was calculated by averaging the relative increase in biomass at the four consecutive time points of the maxima of ammonia concentration. We have chosen the value of the conversion factor *k* such that the doubling time is in agreement with the experimental values^29^.

### Model assumptions

- *G*_*E*_ is the glutamate obtained from the environment and is in a large excess, hence considered constant.
- *k*_*2*_*AG*_*p*_ describes the input for biomass production, i.e. for growth.
- As a simplification, we assumed that only the interior cells produce ammonia since that produced by the peripheral cells is rapidly lost to the environment.
- Loss of ammonia due to diffusion is much larger than that taken up by the periphery to produce biomass. Therefore, the term *k*_*2*_*AG*_*p*_ does not appear in equation (1b).
- Uptake of glutamate (*G*_*p*_) is dependent on itself because glutamate represents the total amino acid and thus protein concentration in the biofilm periphery and can be assumed, in rough approximation, to be proportional to the concentration of various transport proteins embedded in the cell membranes. The greater the concentration of these proteins, the higher is the glutamate uptake rate.

## Results

### Steady states

The steady states of the system can be calculated analytically. This gives a trivial steady state:

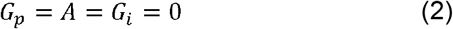

and a non-trivial state:

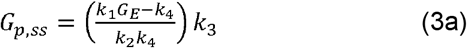

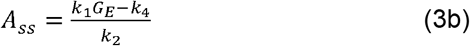

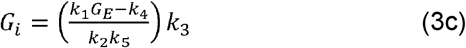

It is worth noting that the concentrations at the latter state are linear functions of *G*_*E*_. The trivial and non-trivial steady states are stable if *k*_*1*_*G*_*E*_ − *k*_*4*_ is negative or positive, respectively^14^. At the threshold, a transcritical bifurcation occurs, that is, the two steady states interchange their stability. At a further threshold, *k*_1_ *G*_E_ = *k*_3_ + *k*_4_ + *k*_5_, the non-trivial, stable steady state turns unstable in a Hopf bifurcation^14^. A Hopf bifurcation is a transition from a stable steady state to limit-cycle oscillations.

### 3.1 Time course shows oscillations

We run the time course calculation of the system (1a-c) for 50 hours with 1000 steps each of size 0.05 hours (3 min). The period for oscillations is about 168 min (2 hours 48 min), with an amplitude of 2.62 mMol for ammonia, 3.71 mMol/mL for interior glutamate and 5.47 mMol/mL for peripheral glutamate (Figure 2). This period is in agreement with the experimental observations^1^, because the parameters have been rescaled accordingly (see above). It can be seen that the three variables oscillate with phase shifts, i.e. asynchronously.

**Figure 2:**
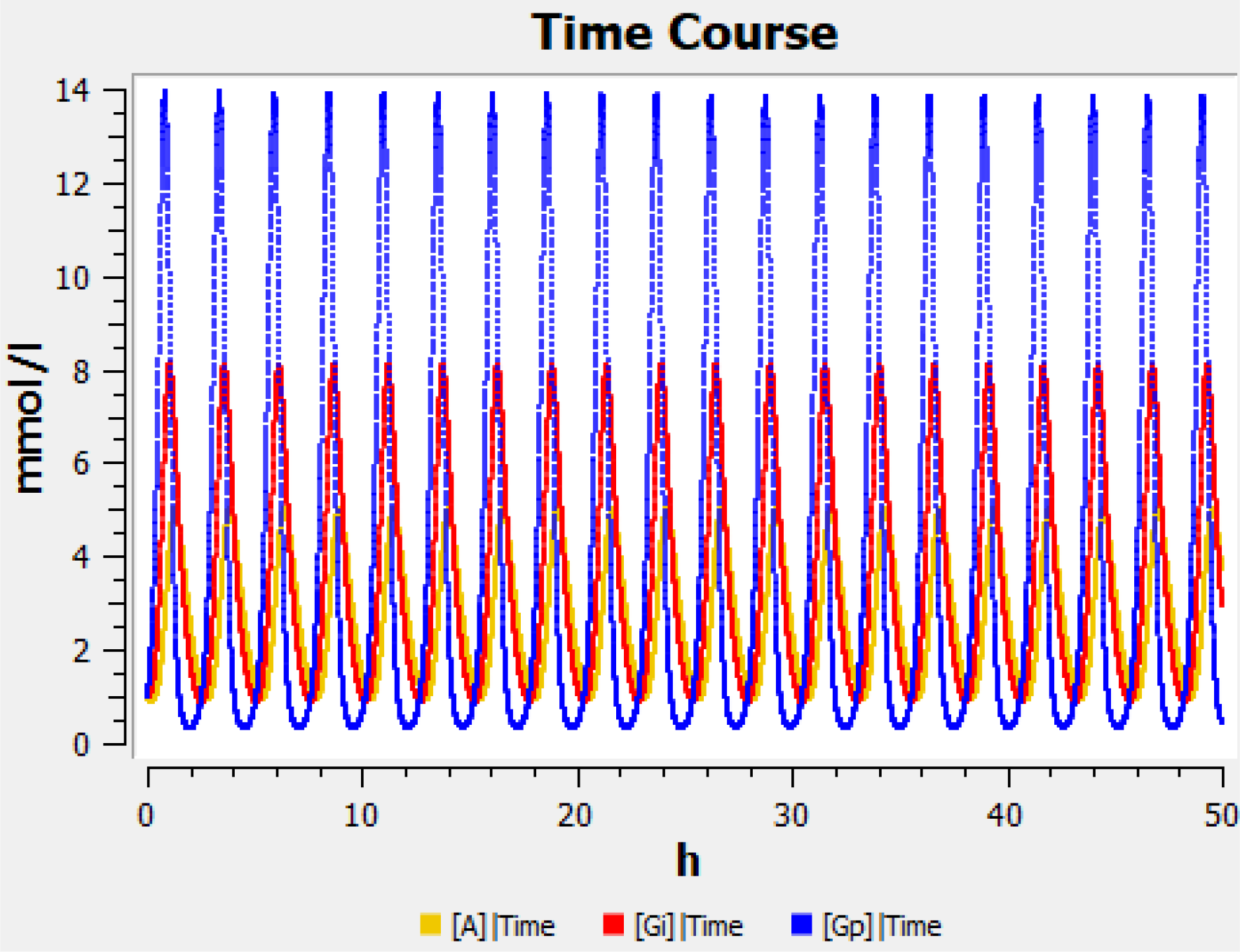
Time course of ammonia (yellow) and interior (red) and peripheral (blue) glutamate as computed by the minimal model. Parameter values: refer to Table 1.

**Table 1:** List of parameters used in the model. These values have been adopted from the publication of Wilhelm and Heinrich^14^ and were rescaled according to the experimental work by Liu *et al.*^1^.

| Parameter | Symbol | Value with unit |
| --- | --- | --- |
| Rate constant of glutamate diffusion from environment to biofilm | $k_1$ | $0.2735 \text{ (mM} \cdot \text{h)}^{-1}$ |
| Biomass formation coefficient | $k_2$ | $2.3 \text{ (mM} \cdot \text{h)}^{-1}$ |
| Rate constant of ammonia diffusion | $k_3$ | $3 \text{ h}^{-1}$ |
| Rate constant of glutamate diffusion within biofilm | $k_4$ | $2 \text{ h}^{-1}$ |
| Ammonia production coefficient | $k_5$ | $2.3 \text{ h}^{-1}$ |
| Glutamate concentration in the environment | $G_E$ | 30 mM |

In order to see the interdependence between the variables of our model, we plot the phase portrait of all three variables for various values of *G*_*E*_ (Figure 3). We found that *G*_*E*_ values below 15 mMol eliminate the oscillations (even damped ones) while values larger than 40 has no qualitative change on the model. *G*_*E*_ is an appropriate bifurcation parameter because the external glutamate concentration can be changed in experiment.

**Figure 3:**
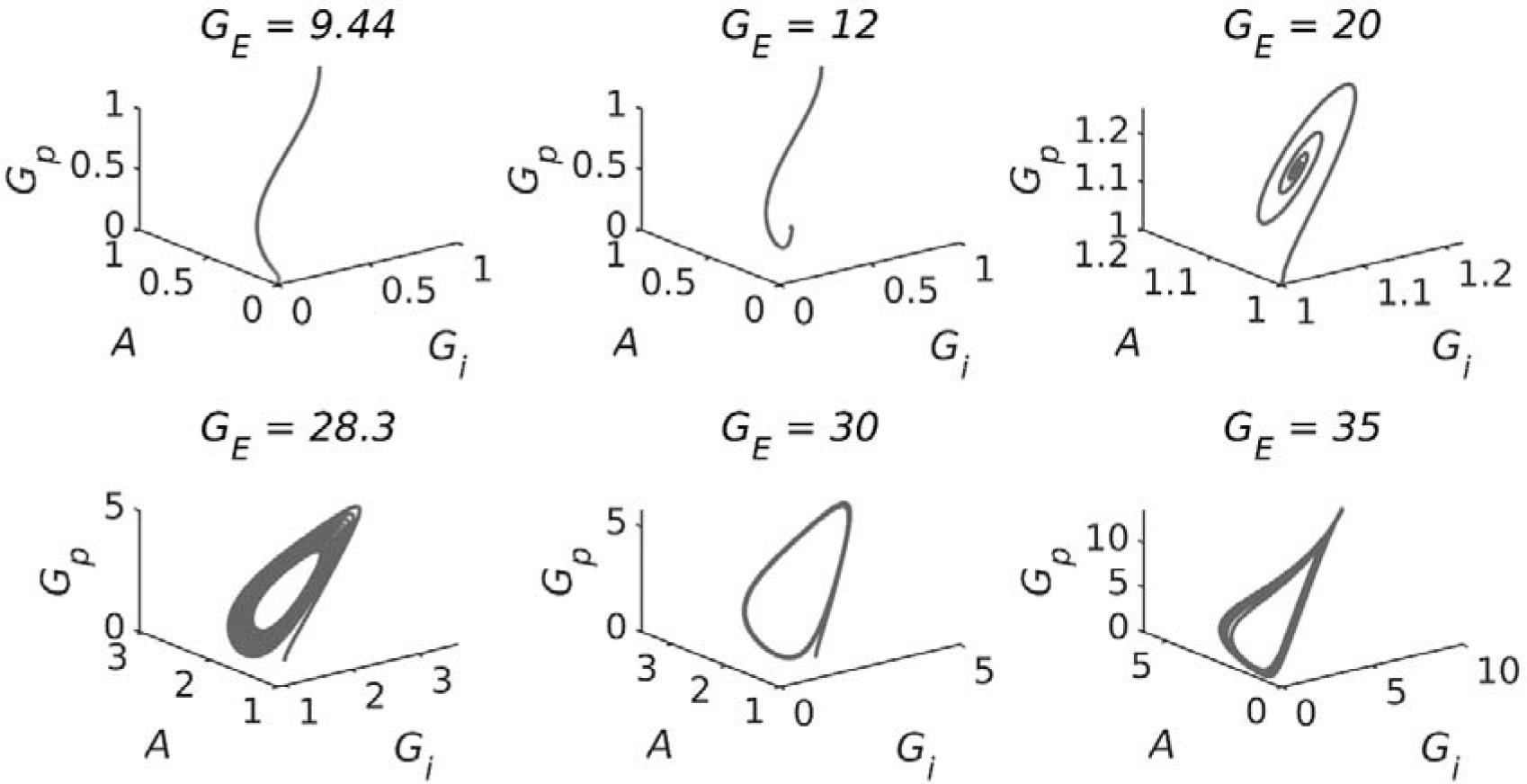
3D phase portrait of all the variables at various values of *G*_*E*_. The trajectory runs anti-clockwise in the perspective shown here. Top three trajectories from left to right depict approaches towards: the trivial SS, the non-trivial steady state (NTSS) in a non-oscillatory way and the NTSS by damped oscillations. The bottom three trajectories depict the convergence towards limit cycles beyond the Hopf bifurcation (*G*_*E*_ = 29.5).

As per the assumptions of our model, *k*_*2*_*AG*_*p*_ is a proxy for the input for the synthesis of biomass from ammonia and glutamate and is thus related to the growth of the biofilm. Biomass production can be described by the following differential equation:

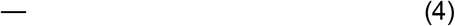

where *k* is a conversion factor and was tuned to 0.1 (mM^2^ h)^−1^. The numerical solution of this equation for various initial values of *G*_*p*_ is shown in Figure 4. It can be seen that there is periodic retardation in growth.

Figure 4 also displays the growth curve in the hypothetical case where *G*_*p*_ and *A* subsisted at steady state (yellow curve). It can be seen that it has a comparable rate of increase as the blue curves.

**Figure 4:**
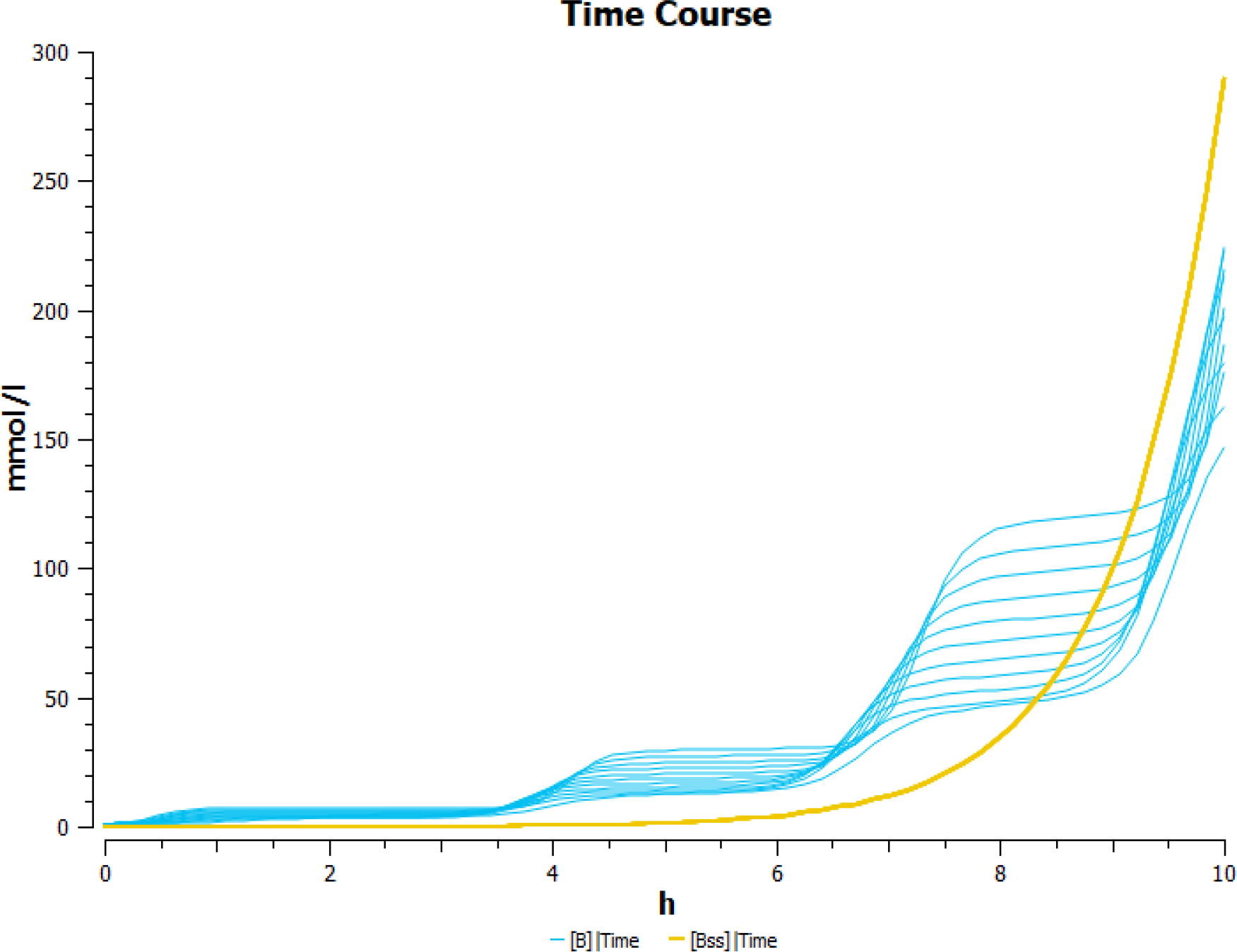
Plot of the time course of growth as calculated from equation (4) for various initial values of *G*_*p*_ from 1 mM – 10 mM (blue curves). On average the curves have a doubling time of about 139 minutes. The yellow curve (Initial value: 0.007 mM) indicates the growth calculated by the steady state values. Parameter values: see Table 1.

The initial value for the growth with constant growth rate (yellow curve) was chosen such that biomass is comparable to that for oscillating growth in the first 10 hours. As can be seen from Fig. 4, it overtakes the oscillating growth at about 9 hours. If the same initial values as for the growth with varying growth rate were chosen, biomass would grow to higher values right from the beginning. Thus, the numerical calculations suggest that oscillating growth is unfavourable for this system.

### Average concentrations and average growth rate

Motivated by the reasoning in the Introduction, we now compare the average concentrations with the steady-state concentrations. As the model under study is a mixture of a Lotka-Volterra equation, equation (1a), and two linear equations (1b,c), it can be assumed that the values equal. To demonstrate this, we divide equation (1a) by *G*_*p*:_
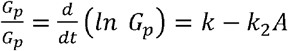 where *k* = *k*_*1*_*G*_*E*_ − *k*_*4*_

We integrate over one oscillation period, *T*:

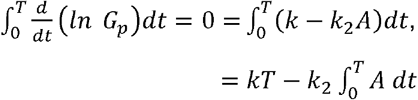

where the integral is zero because *G*_*p*_(*T*) = *G*_*p*_(0)

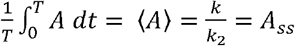

This is, the average *A* equals the steady-state *A*.

Now we calculate the integral of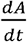:

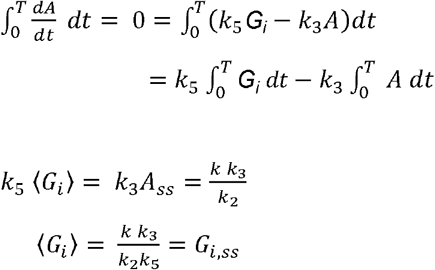

Now, we calculate the integral of 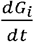 and derive, analogously,

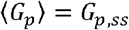

Thus, the average concentration is equal to the steady state concentration.

The question arises whether the oscillations have an effect on the average of the bilinear term *k*_*2*_*AG*_*p*_. This is not immediately clear although Figure 4 suggests that the average value growth is slower than that at equals that of the metabolic steady state. Note that the growth term *kAG*_*p*_*B* is trilinear

To check whether ⟨*AG*_*p*_⟩ = *A*_*ss*·_*G*_*p*,*ss*_, we integrate 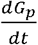 over one period, T:

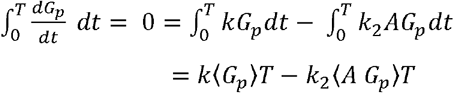

Since⟨*G*_*p*_⟩ = *G*_*p*,*ss*_ and 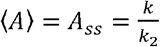, dividing by *k*_2_ and *T gives*

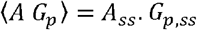

Thus, the average of the bilinear term, which can be interpreted as the input to biomass, is indeed unaffected by oscillations, although the ammonia and peripheral glutamate levels oscillate asynchronously. For two-dimensional Lotka-Volterra systems, this property was shown earlier^18^.

### Bifurcations

Figure 5 shows the two bifurcations: the transcritical bifurcation occurring at *G*_*E*_ = 7.4 mM and the Hopf bifurcation at *G*_*E*_ = 27mM, i.e. the transition from stable steady state to stable limit cycle. These values can also be calculated from the general formulae for the bifurcations ^14^. The steady-state value of *G*_*p*_ is a linear function of the bifurcation parameter *G*_*E*_, as shown in equation (3a). It can be seen that the Hopf bifurcation is supercritical, that is, the amplitude grows gradually starting from zero and the limit cycle is stable right from the beginning.

**Figure 5:**
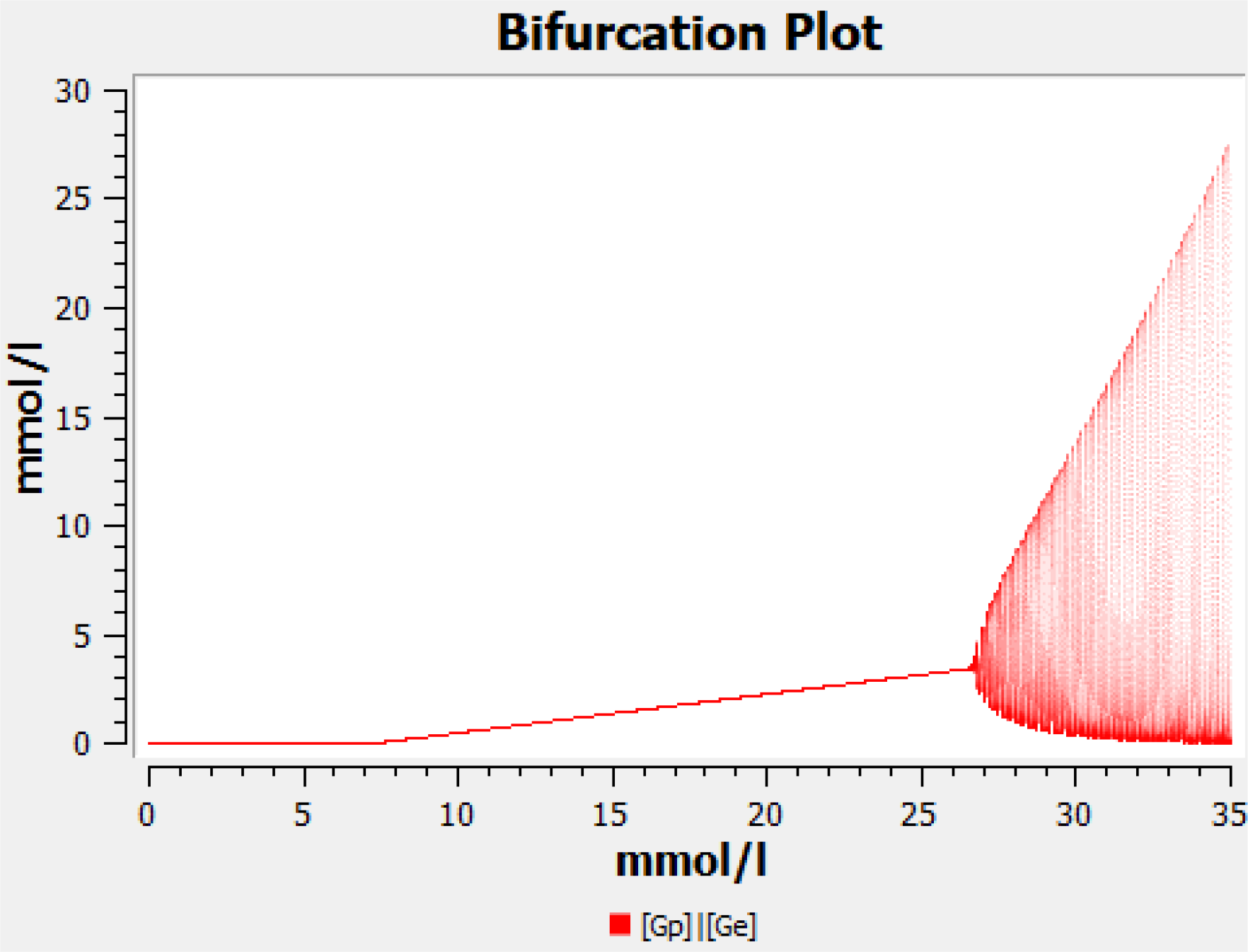
Bifurcation diagram of *G*_*p*_ versus *G*_*E*_. Parameter values are listed in Table 1. The transcritical bifurcation occurs at *G*_*E*_ = 7.4 mMol and the supercritical Hopf bifurcation at *G*_*E*_ = 27mMol. The two arms of the convex hull represent the amplitude of oscillation which widen with increasing value of *G*_*E*_.

Near the Hopf bifurcation, the obtained time course curve (Figure 2) is sinusoidal. For *G*_*E*_ ≫ 27mM the oscillations get spike-like and are no longer sinusoidal. It is of interest to speculate about the physiological advantage of spike-like oscillations. This question has been discussed earlier in the context of calcium oscillations^20,30,31^ Whenever the kinetic effect of the oscillating variable (e.g. in activation of a protein or in a biochemical conversion) is nonlinear and follows a convex function, the spikes contribute more than proportionately to the effect. Thus, spike-like oscillations can lead to a high average effect even at low average value of the variable. In order that oscillations really enable division of labour in the case of biofilms, it can be expected that they should not be sinusoidal. This deserves further studies. A biological explanation of the bifurcations is given in the Discussion.

We checked the parameter sensitivity by varying the reaction constants and found that the model is sensitive to the production of ammonia by the interior (reaction constant *k*_*5*_) (Figure 6). That is, increasing *k*_5_ produces a supercritical Hopf bifurcation (which is laterally inverted) as well and imparts a stabilizing effect and finally the oscillations vanish. Biologically, this agrees with the observation that when the interior produces excess of ammonia, the system no longer oscillates 1.

**Figure 6:**
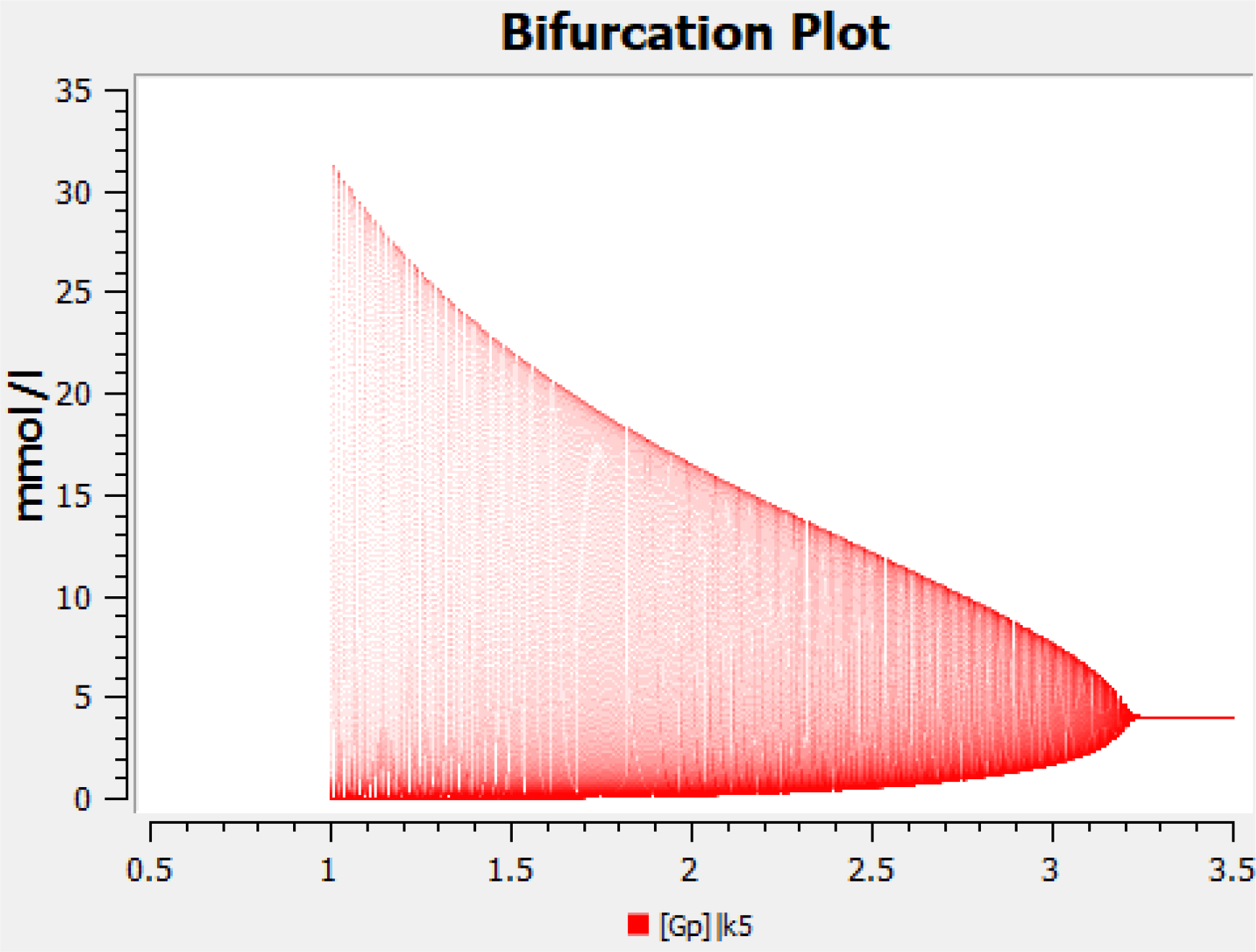
Hopf bifurcation diagram: *G*_*E*_ plotted versus *k*_*5*_. Parameter values are as listed in Table 1 except for *k*_*5*_, which is varied between 1 h^−1^ and 3.5 h^−1^.

### 3.2 Quasi-steady-state approximation

To ascertain the cause of oscillations which could be a positive feedback or a negative feedback or delay, we can study a subsystem by eliminating a variable. This is the Quasi-Steady State Approximation (QSSA). In our system, we see that *G*_*p*_ exerts a positive feedback on itself which is linear and, thus, quite weak. For example, in the Higgins-Selkov oscillator involving two variables, the feedback is quadratic ^11,12^. Moreover, the above system involves a negative feedback: *G*_*p*_ is converted to *G*_*i*_, *G*_*i*_ is converted to *A* and *A* promotes the degradation of *G*_*p*_ (Figure 1). In the Goodwin oscillator, which also consists of three variables, a negative feedback is the cause of oscillation ^32,33^. Inspired by the observation that the oscillations vanish at high *k*_*5*_ values, we applied the quasi-steady-state approximation (QSSA) for *G*_*i*_. This corresponds to the special case where glutamate dehydrogenase is overexpressed and hence the cells produce excessive ammonia ^1^.

We now analyze the special case where reaction k5 is very fast. We can then apply the quasi-steady-state approximation in that the variable *G*_*i*_ attains a quasi-steady state:

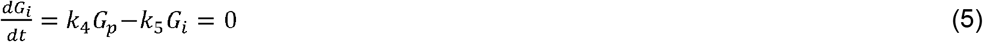

This leads to

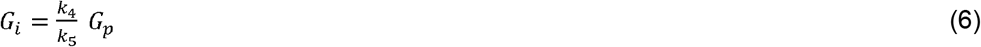

Substituting this into the above model equations (1a, b and c) yields a simplified system:

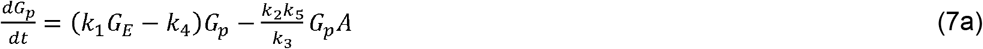

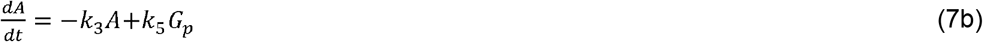

It is of interest to analyze its dynamics. It follows from the general result by Hanusse^34,35^ that it cannot give rise to a limit cycle because it involves two variables and only linear and bilinear terms. However, the question still remains whether it gives rise to a stable or unstable steady state, whether damped oscillations are possible etc.

System (7) shows two steady states:

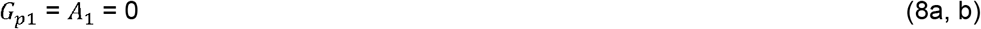

which is the trivial steady state (TSS), and

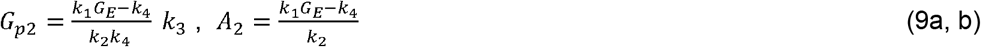

which is the non-trivial steady state (NTSS).

The Jacobian matrix for the NTSS reads:

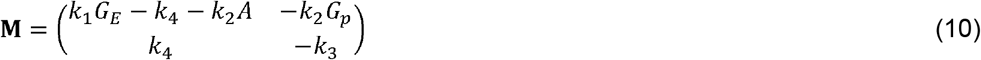

while for the TSS it reads:

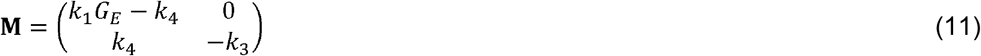

For matrices with such a triangular structure, the eigenvalues are given by the diagonal elements. In our case:

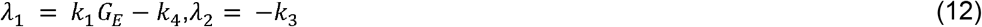

In any case, the eigenvalues are real, so that not even damped oscillations are possible. For *k*_*1*_*G*_*E*_ < *k*_*4*_, both eigenvalues are negative, so that the trivial steady state is a stable node. For *k*_*1*_*G*_*E*_ > *k*_*4*_, one eigenvalue is negative and the other one positive. The steady state then is unstable, it is a saddle point.

For the NTSS (9), the Jacobian matrix becomes:

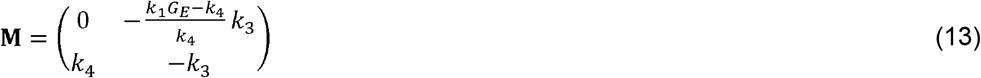

The characteristic equation reads

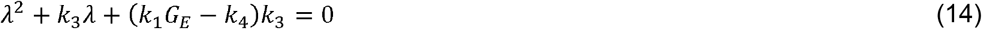

This has the solutions

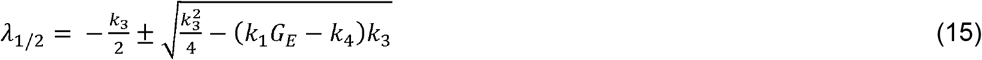

Now, we distinguish three cases:

a. For *k*_*1*_*G*_*E*_ < *k*_*4*_, the term under the square root is positive, so that the root is real. Moreover, it is larger than *k*_3_/2. Thus, one eigenvalue is negative and the other one positive. The steady state then is unstable, it is a saddle point.
b. For 0 < *k*_*1*_*G*_*E*_ − *k*_4_ < *k*_3_/4, the root is again real. It is less than *k*_5_/2, though. Both eigenvalues are negative; the steady state is a stable node.
c. For *k*_*1*_*G*_*E*_ − *k*_4_ > *k*_3_/4, the root is imaginary. Both eigenvalues are complex numbers, with the same negative real part −*k*_3_/2. The steady state is a stable focus. This state is, thus, reached by damped oscillations.

From these calculations, the following conclusions can be drawn. At *k*_*1*_*G*_*E*_ = *k*_*4*_, the two steady states of the simplified system (7) coincide, as in the complete system (1). Since the trivial and nontrivial steady states interchange their stability at that point, it is a transcritical bifurcation.

There is a second transition point where the qualitative behavior changes, at *k*_*1*_*G*_*E*_ − *k*_*4*_ = *k*_3_/4. This is another point than the Hopf bifurcation in the complete system, which is at *k*_*1*_*G*_*E*_ − *k*_4_ = *k*_3_ + *k*_5_. At this transition in the simplified system, a stable node turns into a stable focus. Such a transition must also occur in the complete system between the transcritical bifurcation, where an unstable node turns into a stable node, and the Hopf bifurcation, where a stable focus gets unstable. It is difficult to find it exactly in the 3D system. Beyond this transition, the simplified system shows damped oscillations. This implies *k*_3_ + *k*_5_ that the positive feedback of *G*_*p*_ on itself can be considered as a cause of oscillation, yet not as a cause of a limit cycle. The “inflation” of the oscillation to a limit cycle in the complete system appears to be brought about by the negative feedback via *G*_*i*_. The loop via *G*_*i*_ can, moreover, be interpreted as a delay.

In this subsection, we have considered the special case of high *k*_*5*_. We have proved analytically that the limit cycle disappears in that case. This generalizes the numerical finding shown in figure 6. Thus, to model limit cycle oscillations in biofilms by equation (1), we need the full 3D system with values of *k*_*5*_ that are not too high.

The QSSA for the rate constant *k*_*3*_ is given in the Supplement.

## Discussion

Here, we have used the smallest chemical system showing a Hopf bifurcation to model metabolic oscillations in *B. subtilis* biofilms. That model had been used earlier to describe p53 oscillations^17^. All the terms in the model are linear, except *k*_*2*_*AG*_*p*_, which is bilinear. The model describes metabolic and diffusion processes as outlined above. As an output, the growth of the biofilm (consisting of incremental and halting phases) was also computed (Figure 4).

A major reason of the observed oscillations was demonstrated to be the division of labour between the central and peripheral zones of the biofilm. While the release of usable ammonia is mainly delimited to the former, the production of biomass and, thus, growth, is mainly delimited to the latter.

We have presented a bifurcation diagram, which clearly shows a supercritical Hopf bifurcation (Figure 5). Wilhelm and Heinrich^14,15^ analysed that bifurcation and presented a bifurcation diagram. Here additionally, we show the maxima and minima of oscillations. We analysed the Hopf bifurcation by changing not only the external glutamate *G*_*E*_ but alternatively also the rate constant of ammonia production *k*_*5*_. Interestingly, a recent model^5^ has shown a subcritical bifurcation in describing the behaviour of the stress levels in the biofilm periphery. However, they modelled the stress with a single delay differential equation and did not consider other molecular details, while we do not consider stress. Thus our model is complementary to their model. It is closer to Liu’s original model^1^ but much simpler because it involves only three rather than six variables. We also chose the parameters of the model such that they are in agreement with Liu’s experimental results, namely the period and the amplitude of oscillations.

In our model, peripheral glutamate exerts a positive feedback on itself. Mathematically, this has the form of a bilinear term involving peripheral and external glutamate concentrations. At very low values of *G*_*E*_, the feedback is not strong enough to enable a positive steady state. The system then tends to the trivial steady state. In that state, *G*_*p*_ is zero, so that growth is impossible. Biologically, this can be interpreted in that the biofilm is too small to be viable. This is in agreement with observations in the recent study from the Suel group^5^. At a certain threshold value of *G*_*E*_ (7.4 mM), the non-trivial steady state turns stable in a transcritical bifurcation. Beyond that value, the feedback is strong enough to enable growth. At high values of *G*_*E*_, the feedback becomes so strong that an overshoot occurs: more glutamate is taken up than needed, so that the *G*_*p*_ level transiently exceeds the steady state value. Then, more peripheral glutamate is consumed for release of ammonia or for growth, so that the concentration decreases again. This leads to oscillations.

From a functional point of view, a steady state is quite appropriate^23^. Growth of the biofilm does not require oscillations. However, in this system, oscillations help in mitigating the chemical attack that challenges the biofilm^1^. Furthermore, another study^6^ indicates that oscillations in growth actually help in sharing the nutrients among several biofilms more efficiently. However, not all biofilms show oscillations, indicating that it is not critical for biofilms. Our numerical calculations suggest that the average growth rate is lower as compared to growth at the metabolic steady state.

In contrast, the individual concentration variables (ammonia, peripheral glutamate etc., but not biomass) show the equality property that their average during oscillations equals the value at the unstable state, as usual for Lotka-Volterra systems^19^. Here, we have shown that the bilinear input term *k*_*2*_*AG*_*p*_ exhibits this equality property as well. This may come as a surprise because the ammonia and peripheral glutamate levels oscillate with a phase shift.

In the paper by Liu *et al.*^1^, the oscillations computed by their model have a sinusoidal shape. In our model, such a shape only occurs in the neighbourhood of the Hopf bifurcation. Further beyond it, the shape is more spike-like with the crests being sharper than the troughs.

The question arises whether the model used and analysed here is minimal. On the basis of ordinary differential equation systems (without delays), at least two variables are needed to generate oscillations^9,10^. However, when only linear and bilinear terms are included, at least three variables are needed, as was proved by Hanusse by an analysis of the Jacobian matrix^34,35^. As shown by Wilhelm and Heinrich^14^, such a model requires at least five reactions. Thus, the above model is minimal in terms of the number of variables (criterion with highest priority) and number of reactions, given the type of kinetics. The famous two-variable Brusselator model^36^ and the Higgins-Selkov oscillator involve a cubic term each^11,12^. If the number of reactions is granted the highest priority, the model may look different. Thus, it depends on the criteria what a minimal model is. Note that a delay differential equation^5^ is, from the viewpoint of the number of initial values, of infinite dimension. While our model is not necessarily the simplest, it provides a trade-off between simplicity and adequacy to match the observed oscillation in biofilms.

As for any oscillatory system, it is interesting to elucidate the feedback structure. The term *k*_*1*_*G*_*E*_*G*_*p*_ represents a positive feedback because peripheral glutamate stimulates its own uptake. This is because glutamate is a proxy for the concentration of various transport proteins embedded in the cell membranes. The higher the concentration of these proteins, the higher is the glutamate uptake rate. Since this positive feedback is the driving force for oscillations, at low values of *k*_*1*_*G*_*E*_, we observe a steady state rather than oscillations. In glycolytic and calcium oscillations, the cause of oscillations is also a positive feedback^9,10,12,37^ while in a Goodwin oscillator it is a negative feedback^32,33^.

In addition to the positive feedback there is also a negative feedback loop in the system (see figure 1). As seen from the differential equations, peripheral glutamate positively influences internal glutamate which positively affects ammonia, which then negatively influences peripheral glutamate. Thus, the overall effect is inhibitory. This feedback structure of the Wilhelm-Heinrich model has been stressed earlier^17^.

By applying the quasi-steady-state approximation (QSSA), we have proved analytically that the limit cycle disappears if glutamate is degraded very fast or ammonia diffuses very easily. The QSSA was applied here just to study the extreme case of very fast diffusion. This case corresponds to a situation realised in experiments by Liu *et. al.* where they overexpress the glutamate dehydrogenase leading to an excessive production of ammonia^1^. In that situation, no oscillations were observed. In contrast, in the case where oscillations occur, a description by a simple mass-action system requires three variables.

The model analysed here has several pros and cons. In view of the mathematical analysis, its simplicity is certainly an advantage. In view of an adequate description of the biological and biochemical processes involved, the model may appear oversimplified. For example, describing growth by a bilinear term is quite simplistic; usually it is described by saturation kinetics (e.g. Michaelis-Menten kinetics). In addition the assumption that glutamate uptake by the periphery is proportional to the glutamate concentration as explained above, could be refined in future studies. Moreover, diffusion processes are usually reversible. In the above model, we neglected the backward processes in diffusion, which is justified if concentration differences are high.

Many theoretical and experimental studies have been published on glycolytic oscillations^9,11,12,26,38^. However, these oscillations only occur under very special or even artificial conditions. In living cells metabolic oscillations are rare, while being quite frequent in signalling systems^9,10^. The lights of a car are a helpful analogy: The headlights illuminate the street in a permanent way; there is no point in oscillations. In contrast, the side indicators (as signalling device) flash, that is, they emit oscillating light. Interestingly, in the case of biofilms, metabolic oscillations could provide advantages. While the work described here is quite theoretical, we consider it to be an appropriate basis for refined and more sophisticated models of biofilm oscillations.

## Data availability

### Author information

#### Contributions

Stefan Schuster conceived the idea of applying the Wilhelm and Henrich model to the biofilm system. Ravindra Garde interpreted the model variables and parameters biologically, performed modelling and simulation using COPASI and interpreted the results biologically.

Bashar Ibrahim verified the results using Matlab and also produced figures and mathematical analyses. Stefan Schuster performed QSSA and mathematical proof and mathematical interpretation of the results. Ravindra Garde and Stefan Schuster wrote the manuscript and Bashar Ibrahim edited and structured it.

#### Research Ethics

Our research did not require prior ethical assessment.

#### Animal Ethics

Our research did not require prior ethical assessment.

#### Permission to carry out fieldwork

No permissions were required prior to conducting our research.

## Acknowledgements

We acknowledge David Heckel for his valuable inputs in interpreting the biological implications of the results.

