## Supplementary data for "Differential equation based minimal model describing metabolic oscillations in *Bacillus subtilis* biofilms"

**Applying the quasi-steady-state approximation to ammonia**

Just as in case of the quasi-steady state approximation variable *G_i_* (see main text for details), we apply the same approximation to the variable *A*.

$A=\frac{k_{5}}{k_{3}} G_{i}$	(S1)

From equation (1), we derive a reduced system:

$\frac{{dG}_{p}}{dt}{=k}_{1}G_{E}G_{p}-k_{4}G_{p}-{\frac{{k_{2}k}_{5}}{k_{3}} G_{i}G}_{p}$	(S2)

$\frac{dG_{i}}{dt}=k_{4}G_{p} - k_{5}G_{i}$	(S3)

The system shows two steady states:

${G_{p}}_{1}={G_{i}}_{1}=0$	(S4a,b)

${G_{p}}_{2}=\frac{(k_{1}G_{E}-k_{4})k3}{k_{2}k_{4}}$ , ${G_{i}}_{2}=\frac{(k_{1}G_{E}-k_{4})k3}{k_{2}k_{5}}$	(S5a,b)

The Jacobian matrix reads:

$\mathbf{M}=\left( \begin{matrix} k_{1}G_{E}-k_{4}-\frac{k_{2}k_{5}}{k_{3}}G_{i} & -\frac{k_{2}k_{5}}{k_{3}}G_{p} \\ k_{4} & -k_{5} \end{matrix} \right)$	(S6)

For the TSS, it leads to:

$\mathbf{M}=\left( \begin{matrix} k_{1}G_{E}-k_{4} & 0 \\ k_{4} & -k_{5} \end{matrix} \right)$	(S7)

For matrices with such a triangular structure, the eigenvalues are given by the diagonal elements. In our case:

$\lambda_{1} = k_{1}G_{E}-k_{4}$, $\lambda_{2}= -k_{5}$	(S8)

In any case, the eigenvalues are real, so that not even damped oscillations are possible. For *k_1_G_E_* *< k_4_*, both eigenvalues are negative, so that the trivial steady state is a stable node. For *k_1_G_E_ > k_4_*, one eigenvalue is negative and the other one positive. The steady state then is unstable, it is a saddle point.

For the NTSS (S5), the Jacobian matrix becomes:

$\mathbf{M}=\left( \begin{matrix} 0 & -\frac{k_{1}G_{E}-k_{4}}{k_{4}}k_{5} \\ k_{4} & -k_{3} \end{matrix} \right)$	(S9)

The characteristic equation reads:

$\lambda^{2}+k_{3}\lambda+\left( k_{1}G_{E}-k_{4} \right)k_{5}=0$	(S10)

This has the solutions

$\lambda_{1/2}= -\frac{k_{3}}{2}\pm\sqrt{\frac{k_{3}^{2}}{4}-\left( k_{1}G_{E}-k_{4} \right)k_{5}}$	(S11)

Now, we distinguish three cases:

a) For $k_{1}G_{E}$ < *k*_4_, the term under the square root is positive, so that the root is real. Moreover, it is larger than *k*_3_/2. Thus, one eigenvalue is negative and the other one positive. The steady state then is unstable, it is a saddle point.
b) For 0 < $k_{1}G_{E}$ – *k*_4_ < $\frac{k_{3}^{2}}{4k_{5}}$, the root is again real. It is less than *k*_3_/2, though. Both eigenvalues are negative; the steady state is a stable node.
c) For $k_{1}G_{E}$ – *k*_4_ > $\frac{k_{3}^{2}}{4k_{5}}$, the root is imaginary. Both eigenvalues are complex numbers, with the same negative real part −*k*_3_/2. The steady state is a stable focus. This state is, thus, reached by damped oscillations.

The results of this approximation are similar to those in the main text section on QSSA. The transition between stable node and stable focus occurs at $k_{1}G_{E}$ – *k*_4_ = $\frac{k_{3}^{2}}{4k_{5}}$.
